# Modifications in the piperazine ring of nucleozin affects anti-influenza activity

**DOI:** 10.1101/2022.10.20.513012

**Authors:** Erick Correa-Padilla, Alejandro Hernández-Cano, Gabriel Cuevas, Yunuen Acevedo-Betancur, Fernando Esquivel-Guadarrama, Abraham Madariaga-Mazon, Karina Martinez-Mayorga

## Abstract

The infection caused by influenza virus is a latent tret, to contribute on the advancement of the discovery and design of nucleozin analogs, a molecule with antiinfluenza activity, we analyzed nucleozin analogs with modifications in the piperazine system, which leads to molecules with larger conformational freedom. Following a new nucleozin synthetic strategy, we obtained three new nucleozin analogs, and two of them were biologically evaluated *in vitro* and were less active than nucleozin. The loss of activity in the more flexible molecules highlights the need for the piperazine ring to maintain the activity of nucleozin analogues. Interestingly, this coincides with a QSAR model developed here for the prediction of the anti-influenza activity. The proposed model, along with the synthetic route, will be useful for further development of nucleozin analogues with antiviral activity.

## INTRODUCTION

Influenza is a highly contagious viral infection caused by single stranded negative-sense RNA virus, belonging to the *Orthomyxoviridae* family. This infection affects the respiratory system of the host, affecting the nasal pharyngeal mucosa, bronchi, and pulmonary alveoli. The symptoms of influenza are like those of the common cold; however, influenza can be deadly, especially in vulnerable groups. The mutation rate of the influenza virus; the high frequency of genetic rearrangement; and the antigenic changes in viral glycoproteins, challenge the control of infections, causing even zoonotic interactions, as the avian influenza (H7 and H9) and the swine influenza (Cal/09), with high potential for a pandemic threat. Pharmacological therapy is available. Rimantadine is used to treat or prevent the infection of seasonal influenza (Influenza B) and oseltamivir, rimantivir, zimanavir are preferred for epidemic or pandemic influenza (influenza A).^[1]^ However, emergent strains are potential life-threatening illnesses. Therefore, it is paramount to foster the discovery and development of new therapeutic antiviral agents. Efforts on that direction led to the discovery of nucleozin (**nlz**), a potent inhibitor of influenza A virus infections in *in vitro* and *in vivo* assays ^[2, 3]^. **Nlz**, a piperazine amide, and the corresponding analogues, induce the aggregation of nucleoprotein (**NP**), a protein that plays an essential role on the virus replication cycle. Three-dimensional structures, obtained by X-ray crystallography, of **nlz** and the analogue named **Gerritz 3** bound to **NP**, have been reported in the literature, **PDB ID**: 3RO5^[4]^ and **PDB ID**: 5B7B^[5]^, respectively. In the case of **Gerritz 3**, the three-dimensional structure shows a dimeric complex, formed by two **NP** monomers bridged symmetrically by two molecules of compound **Gerritz 3** [2(**Gerritz 3)**:2**NP**]. In turn, the complex with **nlz** is formed by six **NP** monomers and two **nlz** molecules (2**nlz**:6**NP**). These complexes precipitate in the nucleus of the host cell and do not migrate to cytoplasm where it is necessary for the formation of ribonucleoprotein (**RNP**) and the subsequent assembly of viral structures. Notably, **nlz** is an influenza inhibitor (IC_50_=0.06 μM)^[6]^, more potent than oseltamivir (IC_50_=1 – 10 μM), an approved drug for treatment of influenza infections^[2]^. In addition, the median toxic concentration (TC_50_) of **nlz** and analogs is greater than 250 μM, thus **nlz** and analogues have a wide therapeutic window, thus they are good candidates for further development. To contribute to this area, here we report a novel synthesis of three **nlz** analogues and we also report the *in vitro* evaluation of two of them. In addition, considering these molecules and **nlz** analogues reported in the literature, we present the development of predictive models of activity.

## METHODS

### QSAR models

The workflow to develop predictive models of **nlz** analogues with antiviral activity, is shown in **Figure 1** and consisted of the following steps.

**Figure 1.**
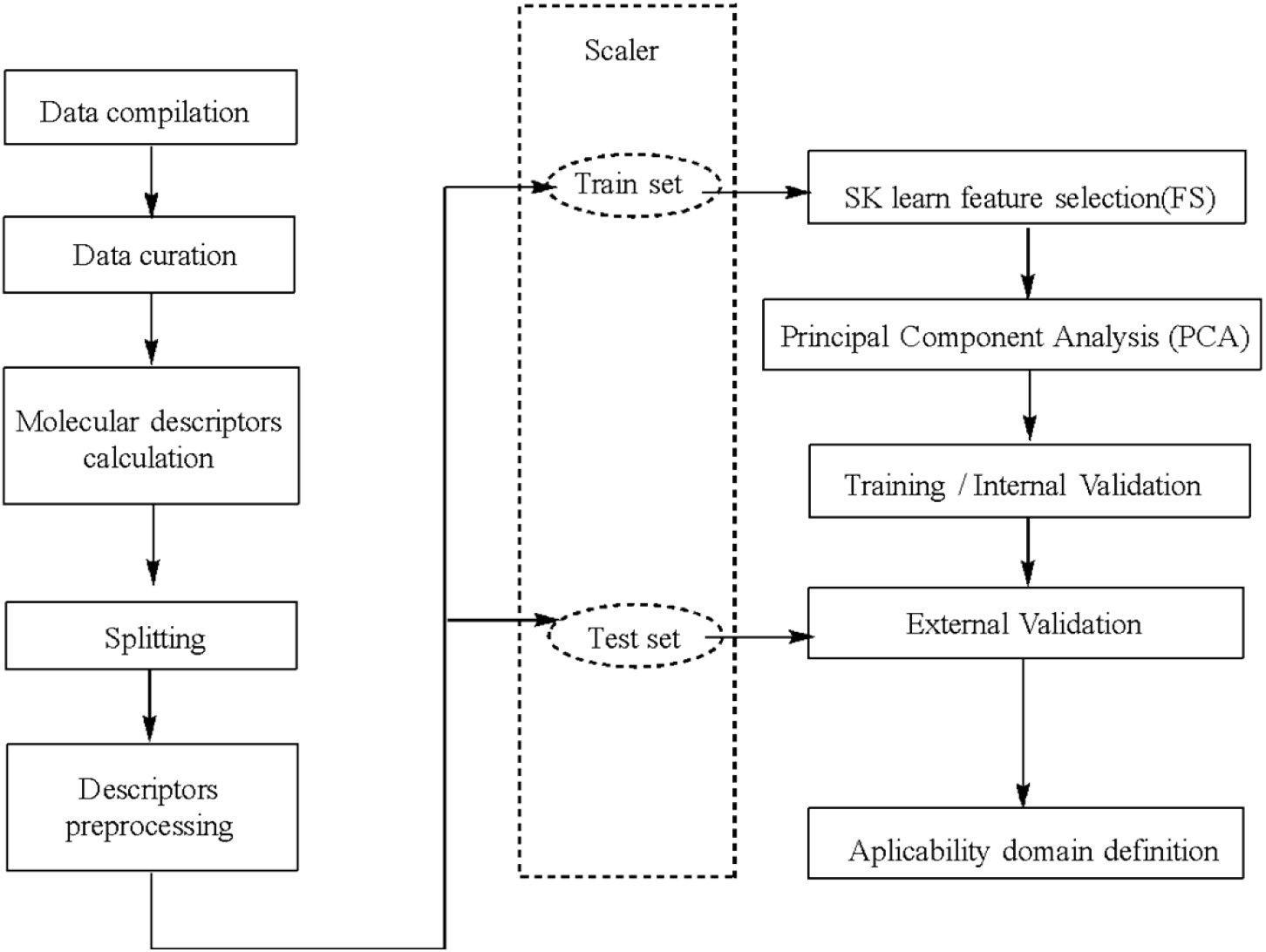
Workflow of the QSAR model.

### Preprocessing

#### Data compilation

The structures of **nlz** analogues were collected from the literature using the keyword nucleozin in Scifinder. The noncurated database consisted of 105 molecules, after removal of datapoints without activity values, a final set contained 74 molecules. The biological activity reported for all these molecules was evaluated in the Plaque Reduction Assay (PRA) with the A/H1N1/WSN/33 strain ^[4, 6-10]^. The antiviral IC_50_ values were transformed to molar units and then to -log(IC_50_), this data is presented in supporting information in table S1. ChemBioDraw Ultra 13.0^[11]^ was used to build the structures. Protonation states were assigned at pH=7.0, and the structures were energy minimized with FFMM94x forcefield in MOE 2019.10. ^[12, 13]^

#### Molecular descriptors

A total of 4185 2D and 3D molecular descriptors were calculated with the software DRAGON (version 7). ^[14]^ Eight additional descriptors (SLogP, Log S, lip_acc, lip_don, TPSA, Weight, b_rotN, b_rotR) were calculated with MOE 2019.10. and analyzed to assess ADME properties.

#### Feature selection and training

The dataset (.csv file format) containing the molecular descriptors and activity values were imported into a notebook in Google Colaboratory (Colab). ^[15,16]^ The modules NumPy ^[17]^, Matplotlib ^[18]^, Pandas ^[19, 20]^and Sci-kit Learn ^[21]^ were used for data handling, analysis, visualization and for the generation of the supervised machine learning regression models. First, the data was splited into a training and test sets, using a stratified split. The training set contains 55 molecules, and the test set contains 19 molecules. The descriptors in the training set, with variance equal to 0, were removed with the algorithm VarianceThreshold, then, the descriptors were scaled with the STDScaler. Then, 70 descriptors with relevance for the prediction of the activity were filtered using the Feature Selection (FS) algorithm from SciKit Learn, using the meta transformer SelectFromModel, and the RandomForest Regressor as base estimator. The feature selection step was performed next, using Principal Component Analysis (PCA) with 16 Principal Components, and using SVR (Support Vector Regressor) as algorithm of regression for the training.

### Model validation

Goodness-of-fit was measured with the coefficient of determination r^2^. Cross-Validation leave-many-out with 10 folds (CV-LMO, folds=10), q^2^ _CV_LMNO_, and Y-scrambling was performed to evaluate the robustness of the model, lastly, the external validation, q^2^ _ext_, allowed to evaluate the predictive power.

### Synthesis of nlz analogues

The chemical structures and numbering of the **nlz** analogues studied in this work are shown in Figures 2-5 and described in the Results section.

**Figure 2.**
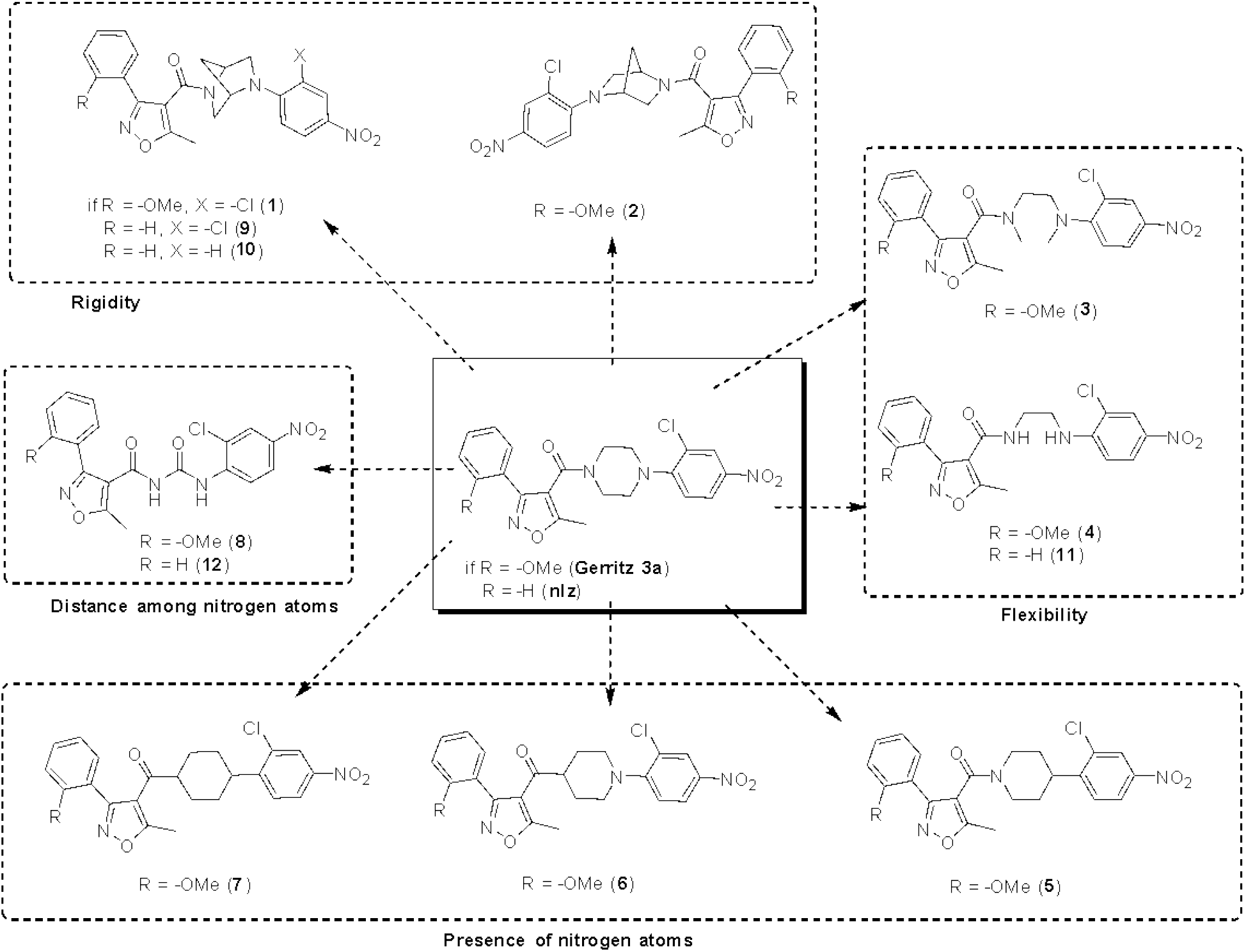
Analogs proposed in this work to study the influence of the piperazine moiety over the antiviral activity within the nucleozin derivatives.

### Biological evaluation

Madin-Darby canine kidney (MDCK) cells were seed at 10^4^ cells per well, in 96-well plates, in Dulbecco’s Modified Eagle’s Medium (DMEM) supplemented with 4 mM -L-glutamine, 1 mM sodium pyruvate, 50 U/mL penicillin, 50 µg/mL Streptomycin and 10% fetal calf serum (FCS), and incubated for 24 hr. at 37 °C. The medium was retired, and cells infected for 1 hr a 37 °C with the influenza A virus strain A/Caledonia H1N1, at a concentration of 0.001 plaque forming units (PFU) per cell. The inoculum was retired, and different concentrations of **nlz** or their analogues (from 0.001µM to 10 µM), diluted in DMEM without serum, added to the cells, and incubated. After 48 hrs., the supernatants were removed, frozen at -20 °C and then titrated in a new 96-well plate with MDCK and incubated for 1 hr. at 37 °C. The inoculum was retired, DMEM without serum added to the cells and incubated for 72 hrs. at 37 °C. The medium was retired, the cells fixed with 4% paraformaldehyde and permeabilized with methanol-acetone 1:1. Then, an ELISA on the fixed-permeabilized cells was performed, using a mouse monoclonal antibody specific for the internal viral protein M1 (ATCC; HB-64), followed by a secondary polyclonal antibody goat anti-mouse Ig’s conjugated with horseradish peroxidase (HRP) (Jackson). The reaction was developed with the substrate o-phenylenediamine dihydrochloride (Sigma) and read in an ELISA reader (Biotech) at 490 nm. The intensity of the signal correlates with the amount M1 produced during the infection. In all the steps, DMEM without serum contained 1 µg of L-1- tosylamide-2-phenylethyl chloromethyl ketone-treated trypsin (TPCK; Sigma) for activation of the virus infectivity. All tissue culture reagents were from GIBCO.

## RESULTS AND DISCUSSION

**Nlz** analogues shown in **Figure 2** vary in the piperazine ring. Compounds **1** and **2** are analogs with the introduction of the 2,5-diazabicycle[2.2.1]heptane (DBH) system, where compound **1** corresponds to the (1*S*,4*S*) diastereomer and compound **2** corresponds to the (1*R*,4*R*) diastereomer. This system has been introduced in a variety of compounds ^[23]^ with antiparasitic, ^[24]^ antibiotic ^[25]^ and anticarcinogenic activity. ^[26]^ It has been suggested that compounds with the DBH system have a better binding ability, compared to the piperazine analogues, due to the rigidity of the bicycle ring. The diamine system of 2,5-diazabicyclo[2.2.1]heptane is traditionally included in screening libraries as a rigid counterpart of the flexible piperazine ring.^[27-31]^ Compounds **3** and **4** exemplify flexible ligands, by substituting the piperazine ring for an ethylenediamine functional group. To assess the role of the nitrogen atoms in the piperazine, we analyzed compounds **5, 6, 7** and **8**. These molecules allowed us to investigate the relevance of the distance among the nitrogen atoms of the piperazine, by diminishing the number of carbon atoms between them.

### Molecular descriptors

An indication of the ADME profile of the molecules studied here was performed through molecular descriptors, calculated with the software MOE 2019.10 ^[12]^(Supporting information). Figure S1 shows a comparison of SlogP and logS values. The SlogP corresponds to the octanol/water partition coefficient, calculated for ∼7000 structures ^[32]^ and is a measure of the lipophilicity. logS is the log of the aqueous solubility (mol/L) calculated from a linear model. The lipophilicity has a large impact on ADME properties, due to its effect in solubility, plasma protein binding (PPB), metabolic clearance, volume of distribution, enzyme/receptor binding among other pharmacological properties. In general, as the lipophilicity increases, the solubility decreases. Also, as the logS value increases (more positive) the solubility increases.

Figure S1, shows the SlogP and logS for **nlz** and other commercial antivirals used for the treatment of the influenza infections, such as oseltamivir, zanamivir, amantadine and rimantadine. Broad spectrum antivirals such as arbidol and acyclovir, foscarnet, peramivir, remdesivir, nirmaltrevir or ritonavir, are also shown. To note, naproxen, acetaminophen and chlorpheniramine are other drugs used in combination with antivirals for the treatment of influenza infections.

The calculated values of SlogP for **nlz** and **Gerritz 3** are 4.17 and 4.18, respectively. These are higher values compared to those of commercial anti-flu drugs, amantadine, rimantadine and oseltamivir, the difference between **nlz** and zanamivir is even larger. The value of **nlz** is also greater compared to those for naproxen, acetaminophen and chlorpheniramine. Not surprisingly, **nlz** and **Gerritz 3** have low aqueous solubility compared with rimantadine, amantadine and oseltamivir, zanamivir. **Nlz** is also less soluble than naproxen, acetaminophen and chlorpheniramine. Thus, although the high lipophilicity is the molecular characteristic that allowed the ligand to interact with the recognition site, this can also be undesirable in terms of pharmacokinetics and ADME properties, due to low solubility. However, arbidol and acyclovir, broad spectrum antivirals, have a higher lipophilicity than **nlz** and **Gerritz 3** and have low solubility values as well.

Figure S2 shows a comparison between SlogP and logS for relevant modifications in **nlz** structure. In the figure S3 and figure S4 is shown the chemical structures for the analogues mentioned above. Modifications in isoxazole-4-caboxamide moiety have been reported in the literature, derivatives of 1*H*-1,2,3-triazole-4-carboxamide are more soluble than **nlz**, even more than the 1*H*-pyrazol-4-carboxamides and phenyl-carboxamides derivatives. In an analogue reported by Liao *et al*, ^[9]^ the absence of the isoxazol ring makes it less lipophilic and more aqueous soluble. In turn, changing the nitrogen atom of the aniline group for a methine group decreases the solubility and increases the lipophilicity.^[10]^

Figure S5 shows that compounds **1** and **2** are less soluble (logS = -6.57) than **nlz** (logS = -6.18) due to the introduction of the methylene bridge, compounds **3** and **4** were slightly more soluble (logS=-6.08 and -6.05, respectively). When the nitrogen atoms are removed in the piperazine moiety the value of logS diminished (logS=-6.61 for ligand **5** and - 6.65 for compound **6**), the enolic tautomeric form of the compound **6** is more soluble than **nlz** (logS = -6.11) and for the case of the compound **7** are even less soluble with logS = -7.66 for the keto form and -7.12 for the enol form. Even with NH groups, compound **8** is less soluble (logS=-6.51) than **nlz**. Thus, the nitrogen atoms have an important contribution for solubility.

### QSAR models

The distribution of IC_50_ antiviral activity values of our curated dataset is shown in Figure 6. Activity values ranges from 4.2 to 7.4. To note, the activity values of training and test sets lie within the same range.

In the pre-processing stage, features with low variance (0 or near 0) were removed. Then, we transformed all the remaining features to have equal variance. By scaling the features, one gives equal chance to all the features on their influence on the model. Thus, the standardized data can be informative. We explored three different scalers (data not shown). For our data, the best scaling algorithm was StandardScaler. As a next step, we selected the most relevant features by assessing their importance using the meta-transformer SelectFromModel, alongside the estimators listed in table S2. To limit the number of features, we first explored the best base estimator with the meta-transformer SelectFromModel, then we fine-tuned this base estimator with the hyperparameters listed in table S3. Lastly, the effect of the number of features was evaluated from 20 to 600 (Table S4), the best models were obtained with 60 to 70 features.

The models were further refined with hyperparameter tuning. ^[21]^ The number of trees in the random forest varied from 50 to 5000. Above 600 trees, the statistics of the models were consistently improved. (Table S3) Max_depth, is the maximum depth of the tree. The default value is None, then nodes are expanded until all leaves are pure or until all leaves contain less than min_samples_split samples (the hyperparameter min_samples_split is the minimum number of samples required to split an internal node, the default value is 2). Max_depth was used in a interval between 1 and 150 (table S3). Here, the best statistics were reached at max_depth = 50.

The hyperparameter Min_samples_leaf is the minimum number of samples required to be at a leaf node. A split point at any depth will only be considered if it leaves at least min_samples_leaf training samples in each of the left and right branches. The default value is 1. In the model Min_samples_leaf were tested in an interval between 2 and 20 (Table S3). Here, the best statistics were reached at Min_samples_leaf = 8.

The hyperparameter min_impurity_decrease refers to a float value and the default value es equal to 0.0. A node will be split if this split induces a decrease of the impurity greater than or equal to this value, it took values between 1.0E^-15^ and 2.0 (table S3). Here, the best statistics were reached at 1×10^−6^.

Lastly, warm_start is a hyperparameter with the Booleans value, the default is False, when set to True, reuse the solution of the previous call to fit and add more estimators to the ensemble, otherwise, just fit a whole new forest. Here, the best statistics was reach with True. All the other parameters that were left as default.

The selected 60 or 70 features were further selected using PCA, reasonably good models contained 15 or more PC Table S5. After the preprocessing, models were generated with several regression algorithms, listed in table 1, exploring from linear regression to Random Forest. Models with the best statistics were SVR and the variant NuSVR. Further refinement was performed using hyperparameter tuning.

**Table 1.** Search for the best regressor algorithm for training.

| Model | Regressor | r<sup>2</sup> |
| --- | --- | --- |
| 1 | LinearRegression | 0.7706 |
| 2 | Ridge | 0.7823 |
| 3 | Lasso | 0.1625 |
| 4 | ElasticNet | 0.7556 |
| 5 | BayesianRidge | 0.7685 |
| 6 | SGDRegressor | 0.7542 |
| 7 | KNearestNeighborRegressor | 0.5791 |
| 8 | SVR | 0.9141 |
| 9 | NuSVR | 0.9103 |
| 10 | LinearSVR | 0.7275 |
| 11 | DecisionTreeRegressor | 1.0 |
| 12 | RandomForestRegressor | 0.9112 |
| 13 | MLPRegressor | 0.9405 |

Using SVR a following step consisted of hyperparameter tuning (table 2) The hyperparameters analyzed were C (the regularization parameter), and max_iter. The first one, C is a parameter that determines the strength of the regularization; must be positive, higher values of C correspond to less regularization, in these cases, the model try to fit each individual data point of the training set as best as possible, while with low values of the parameter C the algorithms put more emphasis on to try to adjust to the majority of data points.^[33]^ The default value is 1.0 Here, the best statistics was reach at 2.0 (model 16 in the table 2). The hyperparameter max_iter is a hard limit on iterations within solver, it is an integer positive value or -1 for no limit (default value). For the best model max_iter = 100 000.

**Table 2.**
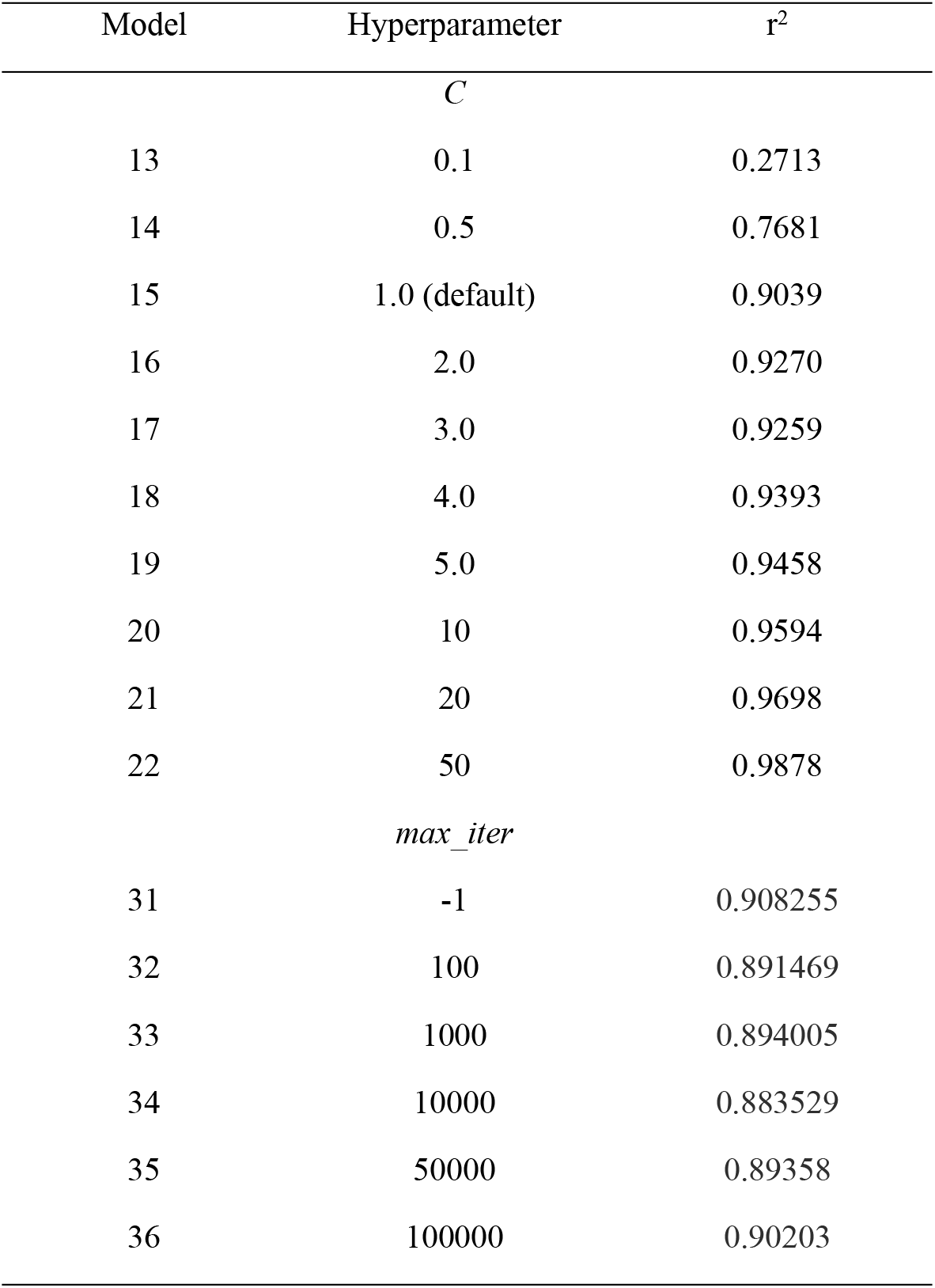
Hyperparameter tuning for SVR regressor.

| Model | Hyperparameter | $r^2$ |
| --- | --- | --- |
| *C* |  |  |
| 13 | 0.1 | 0.2713 |
| 14 | 0.5 | 0.7681 |
| 15 | 1.0 (default) | 0.9039 |
| 16 | 2.0 | 0.9270 |
| 17 | 3.0 | 0.9259 |
| 18 | 4.0 | 0.9393 |
| 19 | 5.0 | 0.9458 |
| 20 | 10 | 0.9594 |
| 21 | 20 | 0.9698 |
| 22 | 50 | 0.9878 |
| *max_iter* |  |  |
| 31 | -1 | 0.908255 |
| 32 | 100 | 0.891469 |
| 33 | 1000 | 0.894005 |
| 34 | 10000 | 0.883529 |
| 35 | 50000 | 0.89358 |
| 36 | 100000 | 0.90203 |

In summary, the best model was developed with SVR, with the hyperparameters C = 2.0, and max_iter = 100 000. The statistics obtained for that model are summarized in Table 3 and the code of Colab is available in the supporting information.

**Table 3.**
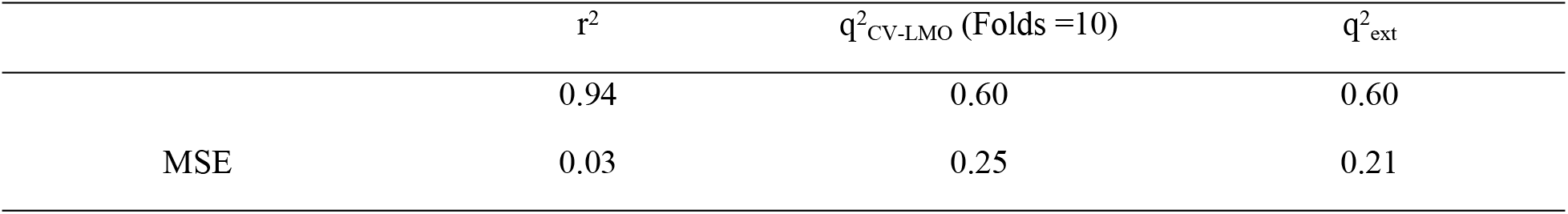
Results for the Support Vector Regressor

| | $r^2$ | $q^2_{\text{CV-LMO}}$ (Folds =10) | $q^2_{\text{ext}}$ |
| --- | --- | --- | --- |
|  | 0.94 | 0.60 | 0.60 |
| MSE | 0.03 | 0.25 | 0.21 |

Figure 7a shows the distribution plot of the values predicted with the SVR model and the experimental values, the train test is shown in blue and the test set is shown in orange, and is defined by a linear correlation. Figure 7b shows the results of the Y-scrambling test with 150 iterations (orange dots), as expected, the q^2^ obtained for the models generated after aleatorization, decreases. Showing that performance of the generated model is not due to chance. Figure 7c shows the applicability domain using the Williams plot. Under that description, the majority of the molecules can be defined by the model. The statistics of the best model are r^2^ = 0.94, q^2^ _ext_= 0.60, and q^2^ = 0.60. The activity predicted for analogues **1** – **12** are shown in Table 4. Interestingly, **Geritz 3** and **nlz** are predicted more actives than the **nlz** analogues, in agreement with the preliminary experimental data presented here.

**Table 4.** Predicted activity for nucleozin analogues.

| Compound | Experimental | Predicted |
| --- | --- | --- |
| Nlz | 7.22 | 6.69 |
| Gerritz 3 | 7.39 | 7.19 |
| compound 1 |  | 5.87 |
| compound 2 |  | 5.86 |
| compound 3 |  | 6.03 |
| compound 4 |  | 5.50 |
| compound 5 |  | 6.60 |
| compound 6 |  | 6.59 |
| compound 6 enol |  | 6.59 |
| compound 7 anti |  | 6.04 |
| compound 7 enol |  | 6.25 |
| compound 7 syn |  | 5.86 |
| compound 8 |  | 5.34 |
| compound 9 |  | 5.54 |
| compound 10 |  | 5.29 |
| compound 11 |  | 4.90 |
| compound 12 |  | 5.06 |

### Synthesis

For this work, a new method for the synthesis of **nlz** was designed. In our lab, the reported synthetic procedure for nlz using 1,2-dichlorobenzen^[6]^ was not reproducible. For the nucleozin synthesis shown in figure 3, the compound **13** was mixed with the secondary amine **14** in DMF at 120 °C, in presence of K_2_CO_3_, for gave the compound **15**, via a nucleophilic aromatic substitution reaction. The subsequent deprotection of the *N*-Boc group with 5.5 equivalents of methane sulfonic acid in dichloromethane yielded a quantitative reaction in five minutes, the compound **16** was obtained. After this deprotection, a solution of the amine **16** was added to a solution of a mixed anhydride, generated *in situ* from the carboxylic acid **17**, treated with pivaloyl chloride in presence of triethylamine, as a base, in dry dichloromethane as the solvent, the consequent addition-elimination reaction gave the **nlz** compound. The same synthetic strategy was employed for the synthesis of compound **9** and **10** (figure 4) for these compounds was used the amine (1*S*,4*S*)-*N*-Boc-2,5-diazabicycle[2.2.1]heptane (**19**), reported previously, instead of the amine **14**. Compound **9** was prepared from 2-chloro-4-nitrofluorobenzene (**13**) and compound **10** from the compound 4-nitrofluorobenzene (**18**).

**Figure 3.**
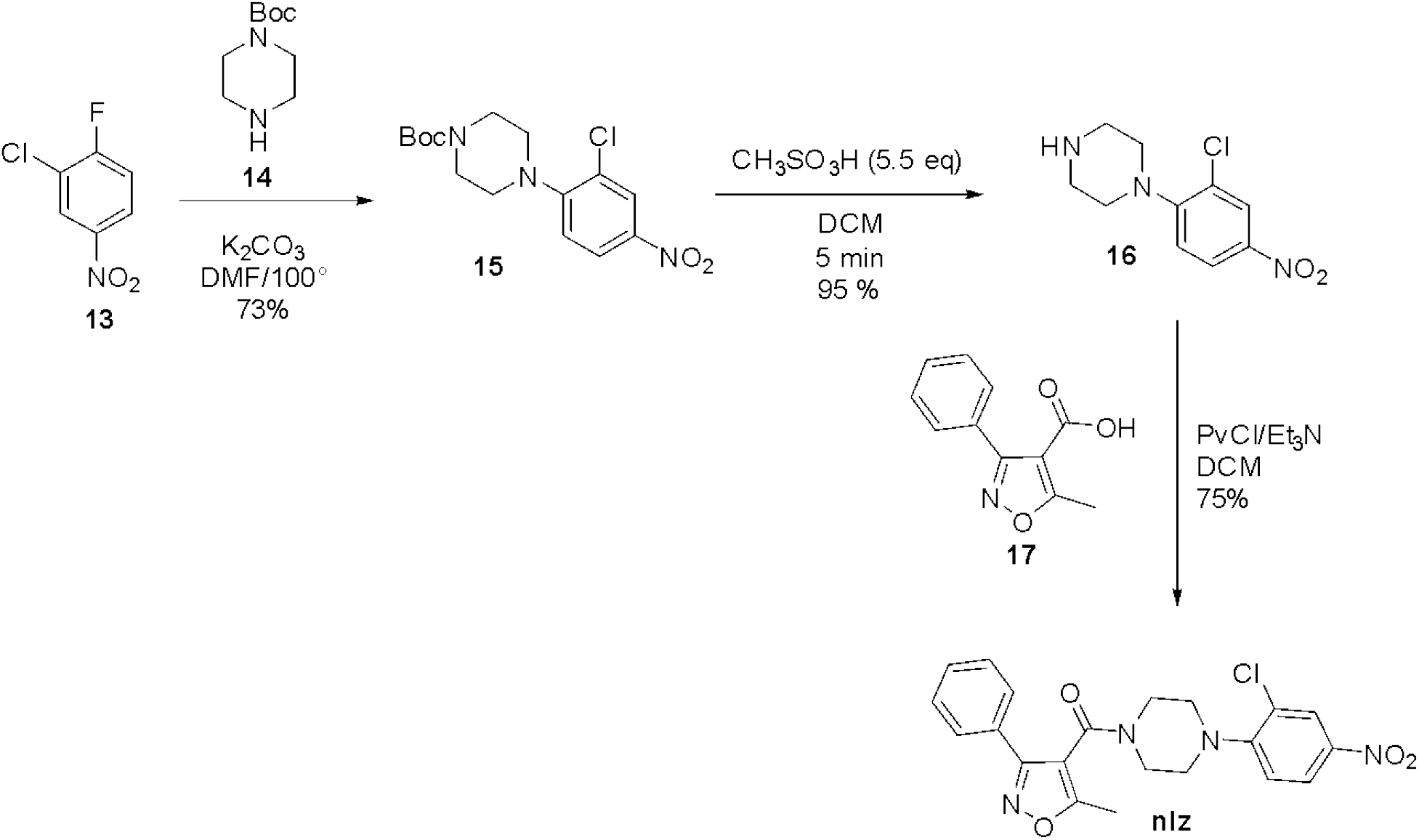
Synthesis of nucleozin.

**Figure 4.**
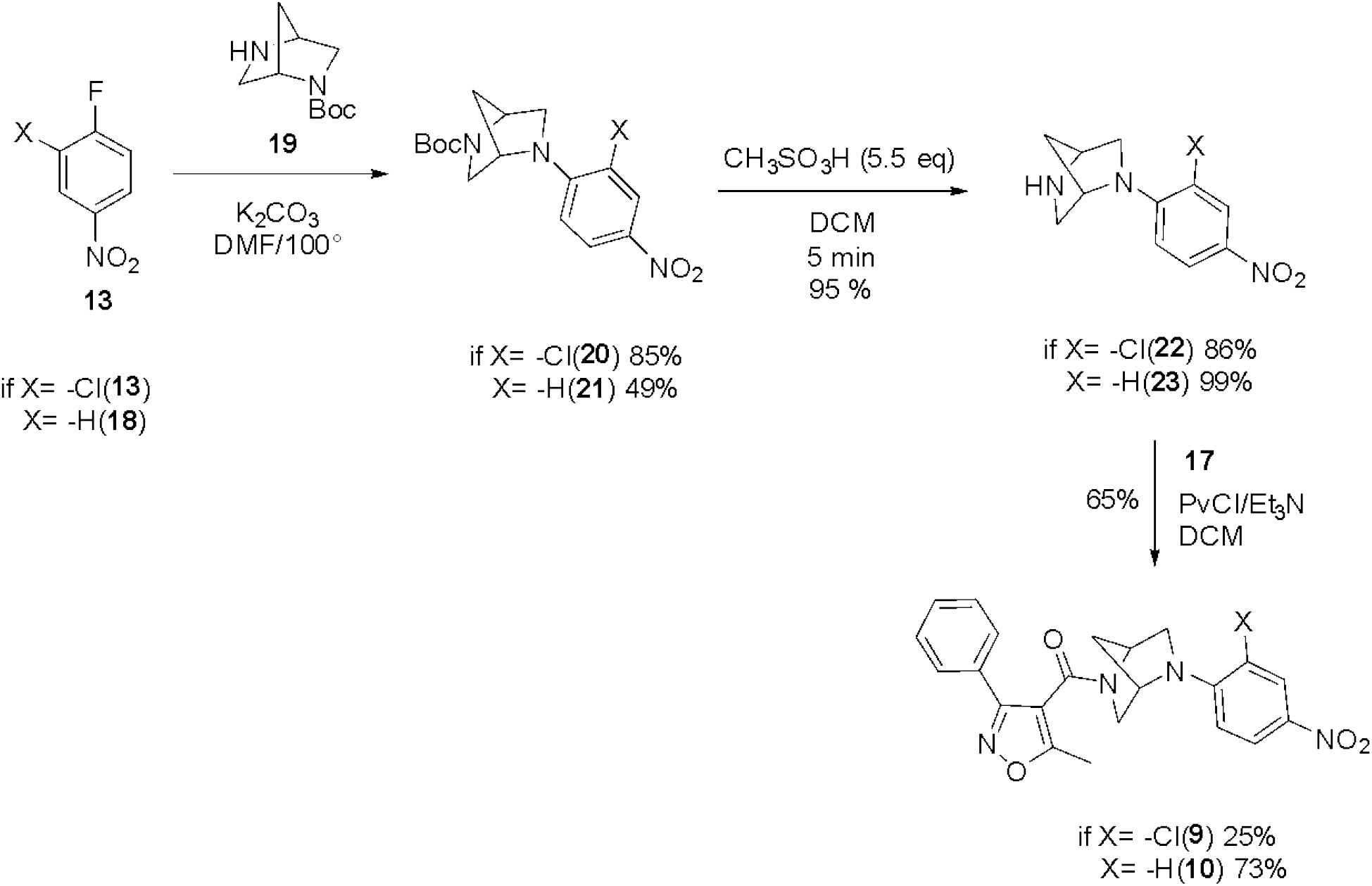
Synthesis of compounds **9** and **10**.

A similar methodology was also used to obtain compound **11**. The ethylenediamine was mono protected using the Boc_2_O reagent to obtain compound **24** which was mixed with an acyl chloride generated *in situ* from the carboxylic acid **17** treated with oxalyl chloride in dry dichloromethane in the presence of triethylamine as a base, and drops of dimethylformamide, rendering compound **25**. Then, for the deprotection of the *N*-Boc group, the compound **25** was treated with methane sulfonic acid, to form the primary amine **26**, followed by a nucleophilic aromatic substitution, using **13** as an electrophile for obtain the compound **11** (figure 5).

**Figure 5.**
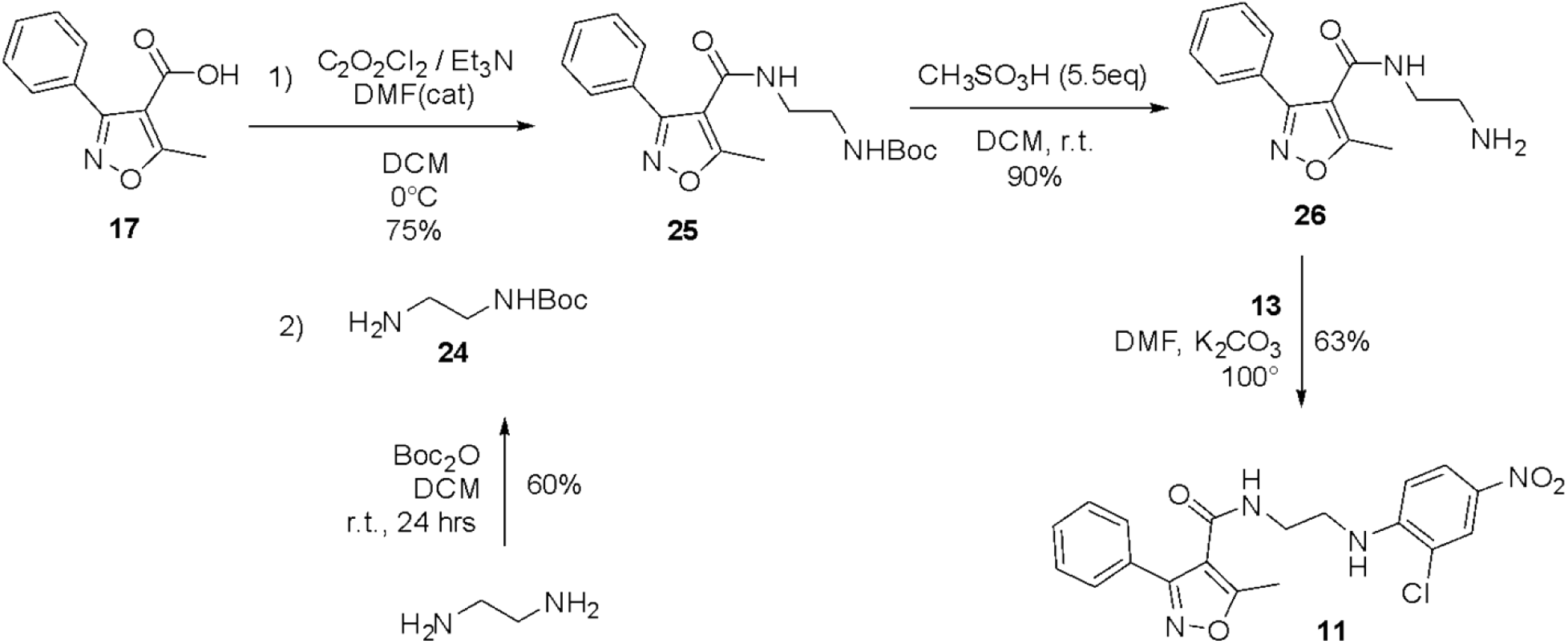
Synthesis of compound **11**.

**Figure 6.**
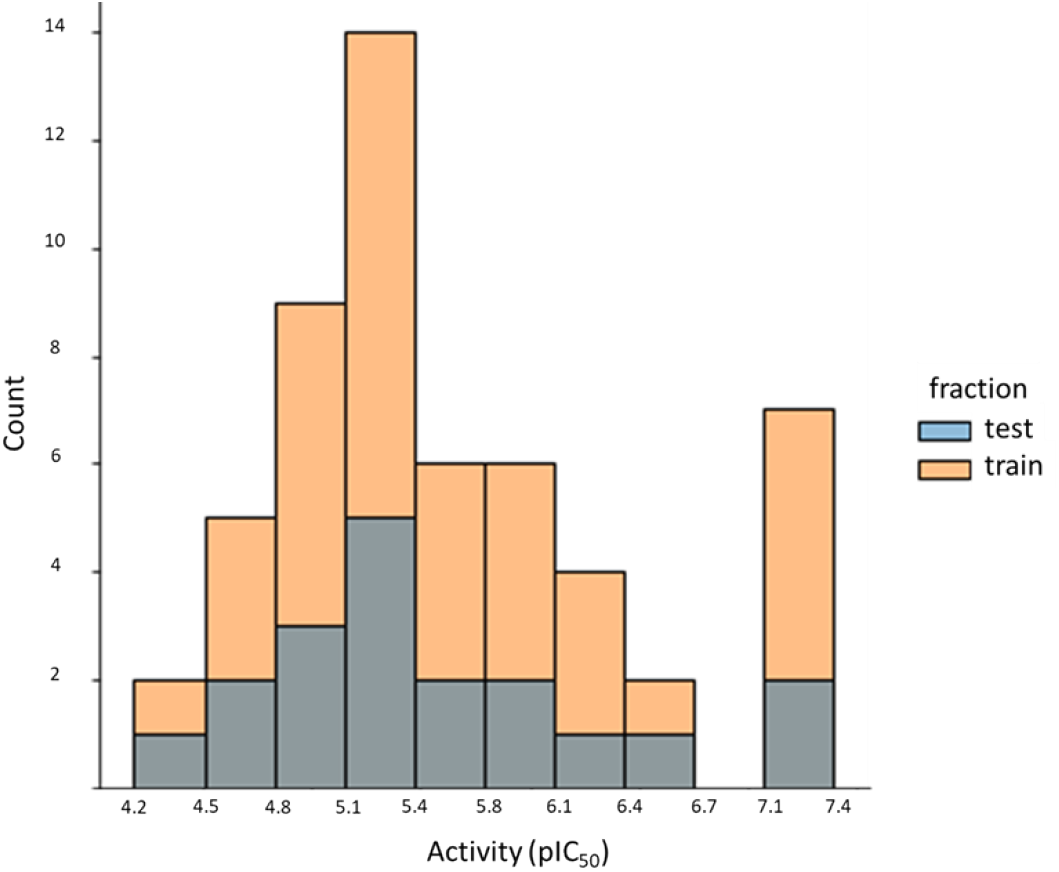
Distribution of pIC50 values for nucleozin derivatives. Molecules in the training set are shown in orange, and molecules in the test set are shown in gray. The inhibitory potency of the test set falls within the interval of pIC50 values of the training set.

**Figure 7.**
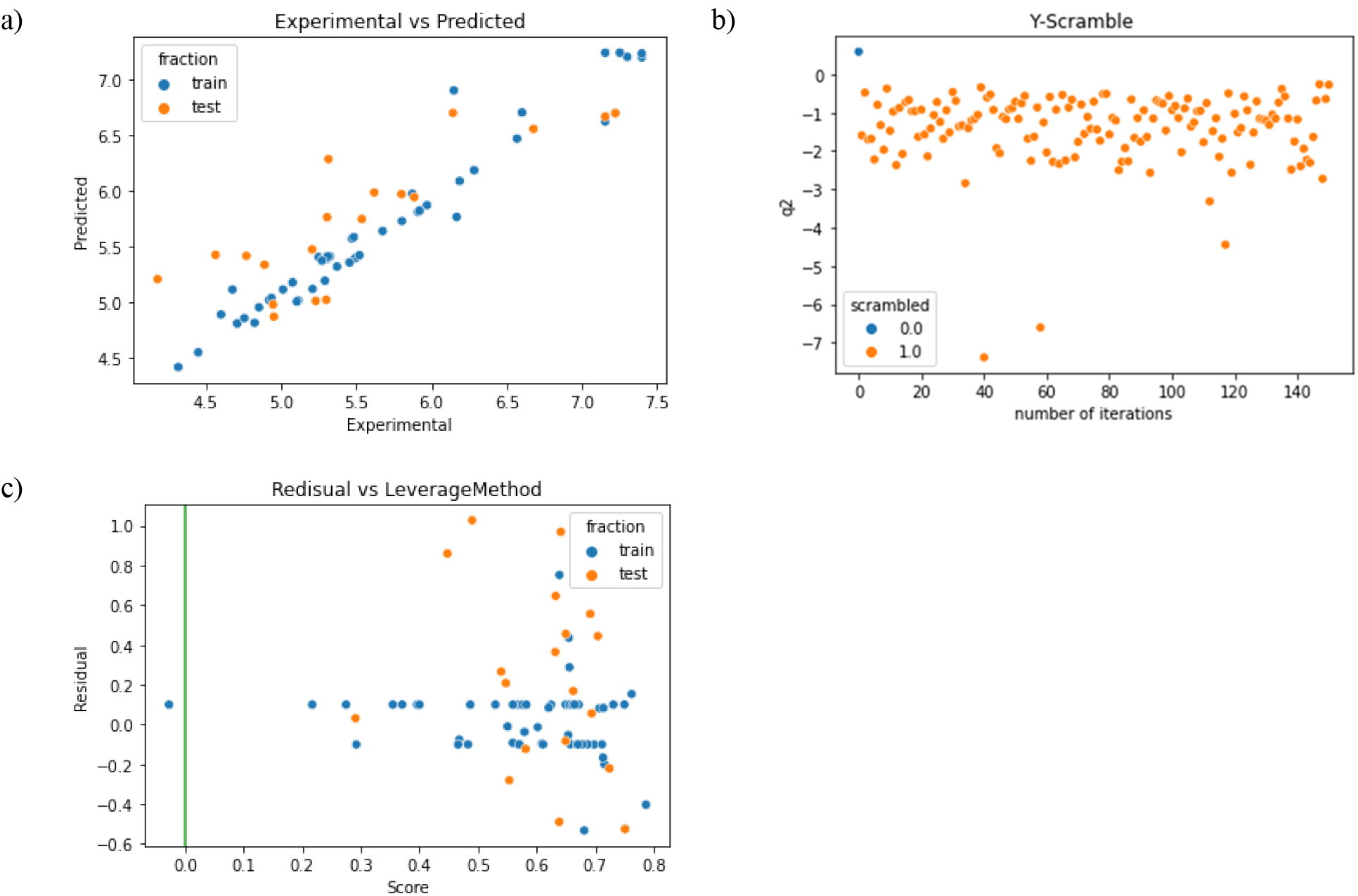
Results of QSAR model. **a)** Regression plot of the SVR model. The plot shows the predicted values by SVR model against the experimental for the train set (in blue) and test set (in orange). **b)** Y-Scramble test for the predicted model: in the plot is shown, in orange, the results of the q^2^ _CV_LMNO_values for randomized models and in blue the q^2^ _CV_LMO_ values for the true model (no randomized). **c)** Williams plot for the calculated model: the Applicability Domain was calculated by the leverage method.

Thus, here we employed a new route using 4-nitrofluroarenes as feedstock, which efficiently led to **nlz** and some derivatives. The aromatic nucleophilic substitution of the fluor atom was performed in dimethylformamide (DMF) as solvent at 120 °C. Some observations in the laboratory indicated that it was not necessary to add DMF to the reaction mixture for synthesis of compound **20**. For this reason, we carried out the reaction grinding in a mortar with a pestle, for 15 minutes, the monoprotected diamine **19** and the compound **13**, both in solid state, in presence of five equivalents of K_2_CO_3_ as a base, without DMF. The reaction was monitored by thin layer chromatography (TLC) comparing the reaction mixture against the previously characterized compound **20**. After the time had passed, most of the raw materials had been transformed. To isolate the product **20**, it was enough to add 3 mL of hot ethyl acetate (AcOEt) to the reaction medium, filter through a small layer of Celite, recrystallize and isolate from mother liquor. By thin layer chromatography a product with a good degree of purity was observed and the melting point coincided with that reported m.p. = 199.7 °C. This tells us that the substitution reaction can be carried out under free-solvent conditions. Future experiments will be carried out to explore this methodology as a potential synthesis route to new **nlz** derivatives.

### Biological evaluation

The virus load starts to decline in the presence of 1μM of **nlz** and is totally inhibited at 10 μM. Under the same conditions but in the presence of the analogues **9** and **10**, viral growth was not inhibited as is shown in figure 8. It is possible that analogues **9** and **10** will have viral inhibitory effect at larger concentrations. These results show that constraining the core structure of **nlz** (analogues **9** and **10**) results in the loss of activity. Interestingly, the QSAR model developed here, predicts the analogues **9** and **10** as less active, consistent with the experimental observation.

**Figure 8.**
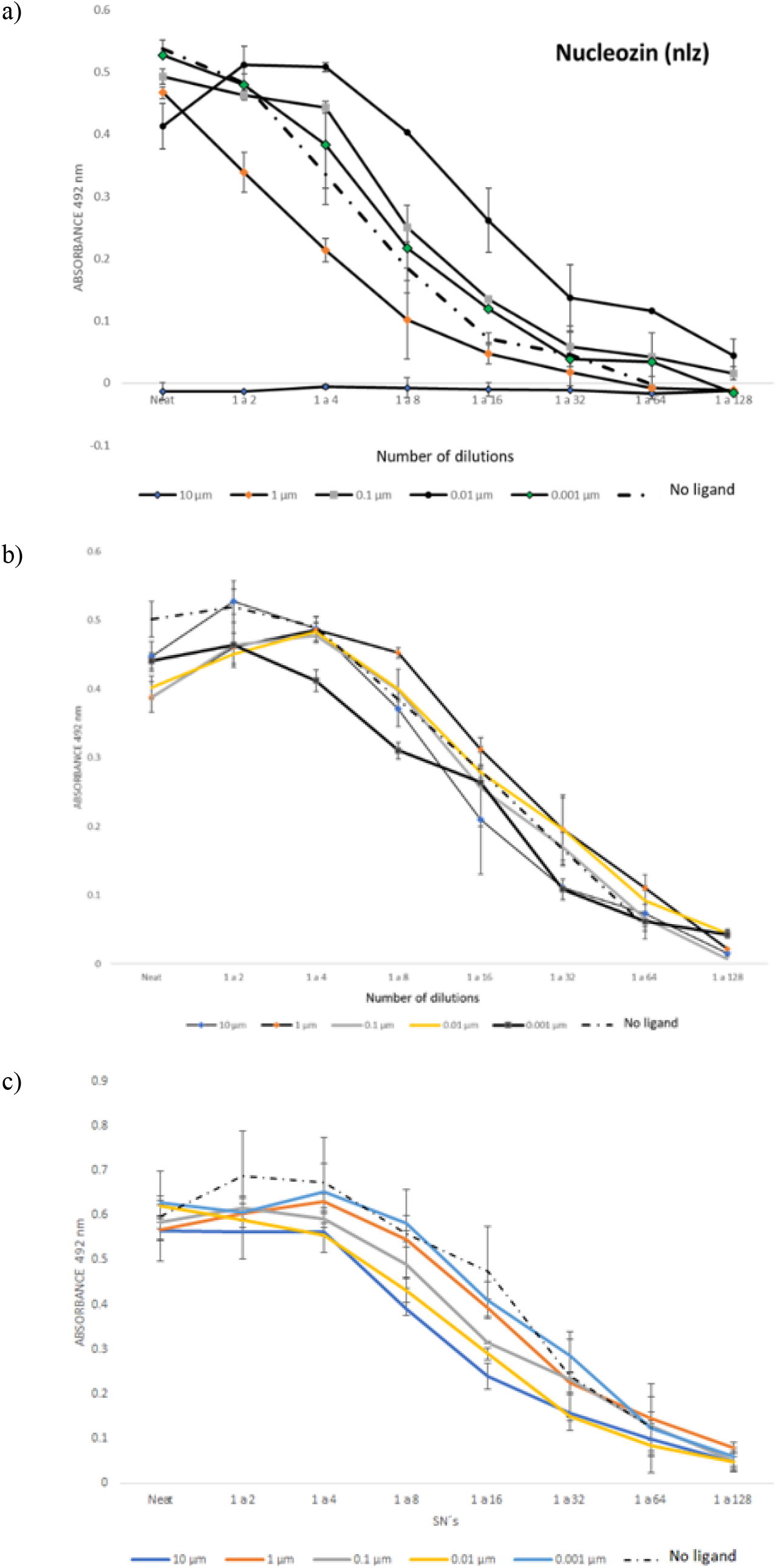
Biological evaluation of **nlz** and analogs in cell cultures of influenza virus-infected MDCK cells. The experiments were performed by infecting MDCK cells with the Influenza virus A/Caledonia H1N1, at a concentration of 0.001 pfu per cell in presence of various concentrations of **nlz** or analogs. After 48 hrs of incubation, the supernatants were obtained and titrated in MDCK cells and incubated for 72 hrs. Cells were fixed-permeabilized and an ELISA on the cells performed, using an anti-influenza M1 monoclonal antibody, followed by a goat anti-mouse Ig–s polyclonal antibody conjugated to HRP. After adding the substrate, the color intensity was read at 492 nm in an automated ELISA plate reader. **a**) Evaluation of **nlz. b**) biological evaluation of the ligand **9. c**) biological evaluation of the ligand **10**.

Additional molecular modeling studies and experimental evaluation of other analogues will provide information on structural modifications required for the improvement of antiviral activity. QSAR models developed here will aid synthetic efforts of new **nlz** analogues.

## Acknowledgements

Eduardo Ruiz-Padilla and Ignacio Regla are gratefully acknowledged for helpful discussions.

